# Polar growth factor PgfA regulates polar peptidoglycan synthesis as well as mycolate synthesis in *Mycobacterium smegmatis*

**DOI:** 10.64898/2026.03.27.714885

**Authors:** Karen E. Tembiwa, Alena M. Truong, Cindy T. Nguyen, Kuldeepkumar R. Gupta, E. Hesper Rego, Cara C. Boutte

**Author notes:** Address correspondence to Cara C. Boutte, E. Hesper Rego.

## Abstract

The mycobacterial cell envelope consists of multiple covalently linked layers that must be synthesized in a coordinated manner to maintain cell wall integrity. Despite the importance of this coordination, its molecular mechanisms remain poorly understood. PgfA (polar growth factor A) interacts with trehalose monomycolate lipids (TMMs) (1) and the TMM transporter MmpL3 (1, 2). PgfA promotes TMM transport in the periplasm and functions as an upstream regulator of polar growth. How TMM transport is linked to the expansion of the entire multi-layered cell wall is unclear. Here, we provide evidence that PgfA regulates peptidoglycan metabolism. We show that PgfA localization correlates with peptidoglycan metabolism and that PgfA can function as both an activator and inhibitor of peptidoglycan metabolism. We further explore the role of TMMs in polar growth and find evidence that periplasmic TMMs are a signaling molecule that may regulate polar peptidoglycan metabolism. Finally, we find an epistatic connection between PgfA overexpression and altered TMM levels that suggests that PgfA and TMMs work in the same pathway to regulate peptidoglycan metabolism. Our data are consistent with a model in which TMM-free PgfA inhibits peptidoglycan metabolism, while TMM-bound PgfA promotes polar peptidoglycan metabolism. This work identifies PgfA as a key protein that coordinates synthesis of the peptidoglycan and mycolic acid envelope layers.

**Importance:** The mycobacterial cell envelope consists of multiple covalently linked layers whose synthesis must be coordinated to maintain cell integrity. Despite decades of research on individual envelope components, the molecular mechanisms coordinating synthesis of different layers remain poorly understood. Here, we identify PgfA as a key regulatory protein that coordinates peptidoglycan and mycolate synthesis in mycobacteria. PgfA has both inhibitory and stimulatory effects on peptidoglycan metabolism, depending on the context. Our findings suggest PgfA may act as a regulator that senses mycolate precursor availability and prevents envelope imbalance when these precursors are limiting. This work provides new insight into how mycobacteria coordinate the synthesis of their complex cell envelope, with implications for better understanding mycobacterial physiology and developing antimycobacterial therapeutics.

## Introduction

*Mycobacterium tuberculosis*, the causative agent of tuberculosis, is an ancient pathogen that is among the deadliest throughout human history and continues to be the leading cause of infectious deaths worldwide (3). The mycobacterial cell envelope is architecturally complex and therapeutically important (4, 5). Mycobacteria have a multi-layered cell wall consisting of peptidoglycan (PG) covalently linked to arabinogalactan (AG), which in turn is decorated with long-chain mycolic acids that form the inner leaflet of the mycolic acid (MA) outer membrane (6–8). Enzymes that build this wall are targets of both first line and second line anti-tubercular antibiotics (3, 9)

Coordination of envelope layer synthesis is a fundamental challenge across bacterial Phyla. The three layers of the mycobacterial wall are assembled by distinct biosynthetic machineries operating at the same locations within the cell. Synthesis and transport to the periplasm of cell wall precursors appears to happen mostly at or near the cell poles (2, 10–19). However, the cell wall enzymes in the periplasm do not appear to all be polarly localized (4, 19–29), even though cell wall labeling experiments indicate that synthesis of the wall is polar (18, 27, 30–33). These observations suggest that spatial co-localization of biosynthetic enzymes alone is insufficient to explain coordination between envelope layers. In both Gram-negative and Gram-positive species, sharing of the lipid anchor between precursors of PG and other cell envelope components helps coordinate assembly of the envelope (34–38). In mycobacteria, the arabinogalactan precursors (39–41) share a lipid anchor with the precursor of PG (42). Thus, the lipid anchor sharing system may coordinate synthesis of the PG and arabinogalactan layers of the wall. Mycolic acid lipids do not use the same anchor (43), and so there is no model to explain how PG and MA synthesis might be coordinated.

Recent work has identified PgfA (MSMEG_0317 in *M. smegmatis*, Rv0227c in *M. tuberculosis*) as an essential protein that localizes to the cell poles and division septum(1). PgfA was found to interact physically with MmpL3(1, 2), the essential TMM flippase that transports TMM across the plasma membrane (16). PgfA also binds TMMs, and promotes the transport of TMMs, possibly by helping bring them from MmpL3 to the outer membrane (1). PgfA’s localization correlates with polar growth, and its overexpression can rescue the old pole elongation defect of a Δ*lamA* mutant(1). However, the deletion of *pgfA* is lethal(1) and leads to disassembly of the cell wall, indicating that PgfA has an essential function in construction or maintenance of the wall. The mechanism behind how PgfA’s interactions with MmpL3 and TMMs affect cell wall metabolism and promote polar growth have remained unclear.

We investigated PgfA function in *Mycobacterium smegmatis*, a fast-growing model mycobacterium with the same cell envelope architecture as *M. tuberculosis*(44). Here, we demonstrate that PgfA acts as a bifunctional regulator of peptidoglycan metabolism. Our findings support a model that PgfA senses periplasmic TMM availability through its capacity to bind TMM and may inhibit PG metabolism when periplasmic TMM levels are low, thereby helping to coordinate PG metabolism with mycolic acid availability. Our results point to a previously undescribed role for PgfA as an inhibitor of PG metabolism and provides new insight into how bacteria coordinate synthesis of architecturally complex cell envelopes.

## Results

### PgfA localization correlates with peptidoglycan metabolism and responds to metabolic state

Previous work shows that PgfA localization correlates with polar growth (1). We expect that peptidoglycan metabolism is a major contributor to growth at the pole. To determine the extent to which PgfA localization correlates with PG metabolism, we analyzed the fluorescence intensity distribution of PgfA-mRFP and HADA in an *M. smegmatis* strain expressing native levels of PgfA-mRFP as the sole copy of *pgfA*. In log phase cells, PgfA-mRFP and HADA intensities showed significant positive correlation at both poles (old pole: Spearman ρ = 0.68, p < 0.0001; new pole: ρ = 0.71, p < 0.0001) and within the midcell region (ρ = 0.64, p < 0.0001), indicating coordinated localization of PgfA with sites of active PG metabolism (Fig. 1A). This result is notable in comparison with similar analyses performed with polar regulators Wag31/DivIVA, PlrA and MmpL3, where we previously observed that the localization of these proteins all correlate with HADA only weakly at the new pole and not at all at the old pole (45, 46). That PgfA’s localization correlates more strongly with growth than these other factors supports the idea that it could be a more direct regulator of PG metabolism.

**Fig. 1.**
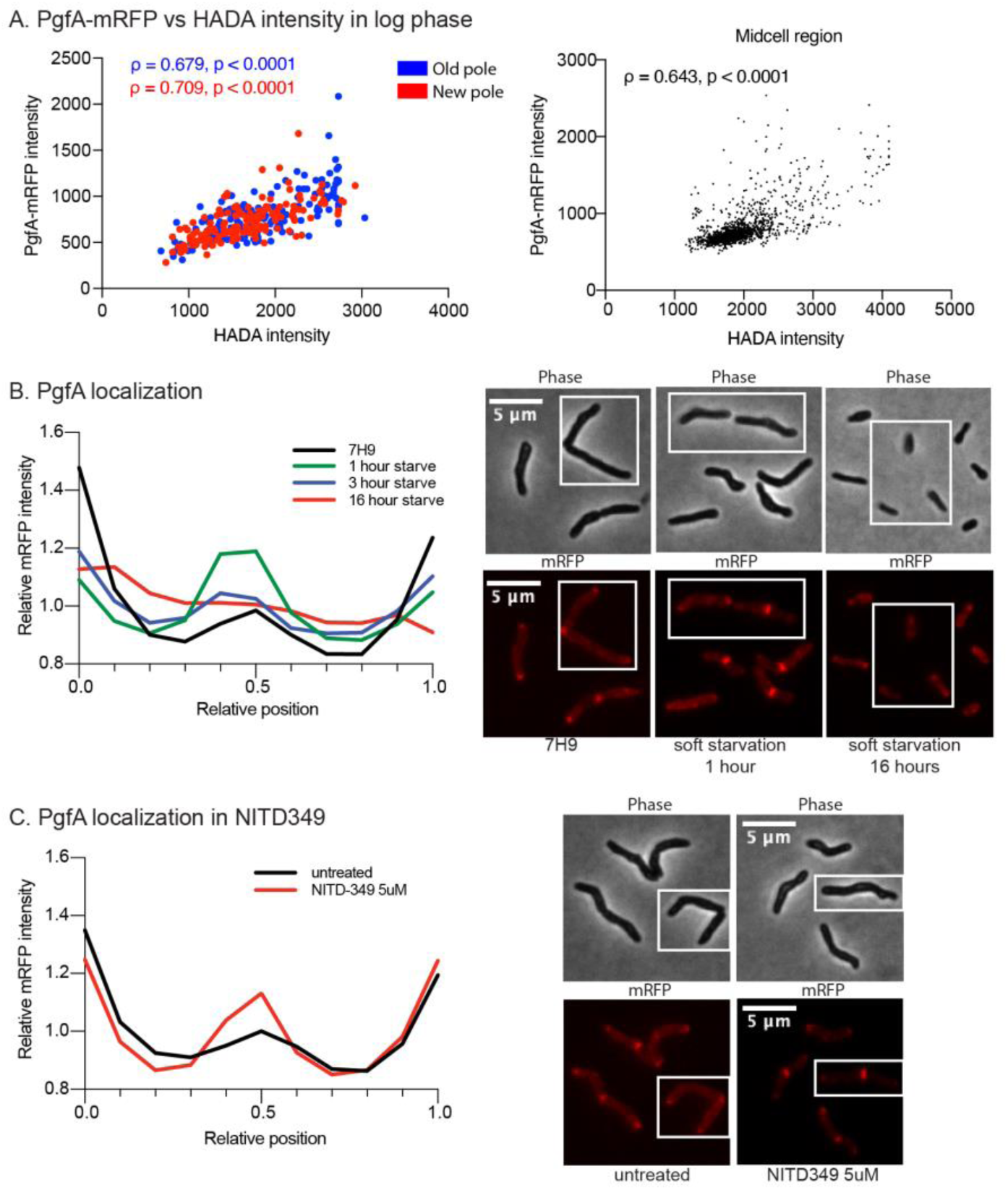
PgfA localization correlates with peptidoglycan metabolism and responds to metabolic state *(A)* Scatter plots of PgfA-mRFP versus HADA fluorescence intensities in log. phase cells. **Left**: polar mean intensities with old pole (blue) and new pole (red) shown separately. **Right**: intensities within the midcell region (normalized positions 0.4-0.6). Spearman rank correlation coefficients (ρ) and p-values are indicated. n = >300 cells. *(B)* Normalized mRFP fluorescence intensity profiles along the cell axis (0 = old pole, 1 = new pole) during progressive nutrient limitation. Cells were grown in 7H9 medium (black) or subjected to starvation in the presence of Tween80 for 1 h (green), 3 h (blue), or 16 h (red). **Left**: Fluorescence intensity profile. **Right**: representative micrographs. Scale bar, 5 μm. All data represents n=>300 cells, >100 per biological replicate. *(C)* Normalized mRFP fluorescence intensity profiles presented as in *(A)*. PgfA localization following 1 h treatment with MmpL3 inhibitor NITD-349 (1.53 µg ml⁻¹). **Left**: Fluorescence intensity profile. **Right**: Representative micrographs. All data represents n=>300 cells, >100 per biological replicate.

Other proteins involved in cell wall synthesis, MmpL3 and PlrA, delocalize from the poles upon starvation (45, 46). To determine if PgfA responds similarly, we examined PgfA-mRFP distribution during progressive nutrient limitation. During starvation in the presence of Tween80, in which cells arrest elongation but reductively divide(47), PgfA initially relocated from cell poles to the division septum within 1-3 hours and became largely delocalized by 16 hours when cell wall metabolism is completely arrested (Fig. 1B). Under complete carbon starvation, delocalization occurred more rapidly, with minimal septal localization at the shorter time points. Prolonged complete starvation (16 h) caused a pattern with large, bright PgfA foci that are excluded from the poles – the significance of this is unclear (Supplementary Fig.1A-B). Correlation analysis across the same growth and stress conditions revealed that the spatial coordination observed in log phase (Fig. 1A) persists during early starvation (1-3 h) but is lost during prolonged severe nutrient limitation (16 h hard starvation; Supplementary Fig. 1C). Together, these results indicate that PgfA localization is highly sensitive to nutrient status and tracks with sites of active cell wall metabolism.

PgfA has been shown to interact with MmpL3 and may be involved in trafficking TMMs across the periplasm (1, 48). To assess how PgfA localization responds to disrupted TMM transport, we analyzed cells treated with NITD-349. Treatment with 1X MIC of NITD-349 caused a modest increase in septal PgfA localization while maintaining strong polar enrichment (Fig. 1C). This contrasts with the dramatic relocalization observed during nutrient starvation. The maintained polar localization under NITD-349 treatment suggests that PgfA localization does not strictly depend on active TMM transport. Since MmpL3 does not delocalize when inhibited(45), the polar PgfA localization may be due to its interaction with MmpL3 at the poles. The relative increase in septal PgfA may represent redistribution as cells shift toward septal division when elongation is impaired, like the pattern observed during early starvation (Fig. 1B).

### PgfA depletion reveals a role in activating PG metabolism at the poles

Our finding that PgfA localization correlates with peptidoglycan metabolism suggests that PgfA may help regulate PG synthesis. Previous work demonstrated that PgfA is essential for survival and its overexpression is necessary for fast polar growth in certain genetic backgrounds(1). To investigate PgfA’s functional role in regulating cell wall synthesis, we used CRISPRi to conditionally deplete *pgfA* in wild-type *M. smegmatis*. Western blot analysis using a custom anti-PgfA antibody confirmed progressive depletion of PgfA protein over the timecourse of depletion (Fig. 2A). PgfA depletion caused progressive changes in cell morphology and distribution of PG metabolism as measured by HADA staining. Phase contrast micrographs show progressively shorter cells beginning at 3h with increasing bulging at later timepoints (Fig. 2B). HADA initially appears to decrease, though not significantly, at 1h, then significantly increased at 3h compared to 1h, before progressively declining through later timepoints (Fig. 2C). This biphasic pattern suggests that PgfA may exist in different states, which have different effects on PG metabolism. Cell length decreased significantly over the course of PgfA depletion (Fig. 2D), confirming that PgfA is essential for cell elongation.

**Fig. 2.**
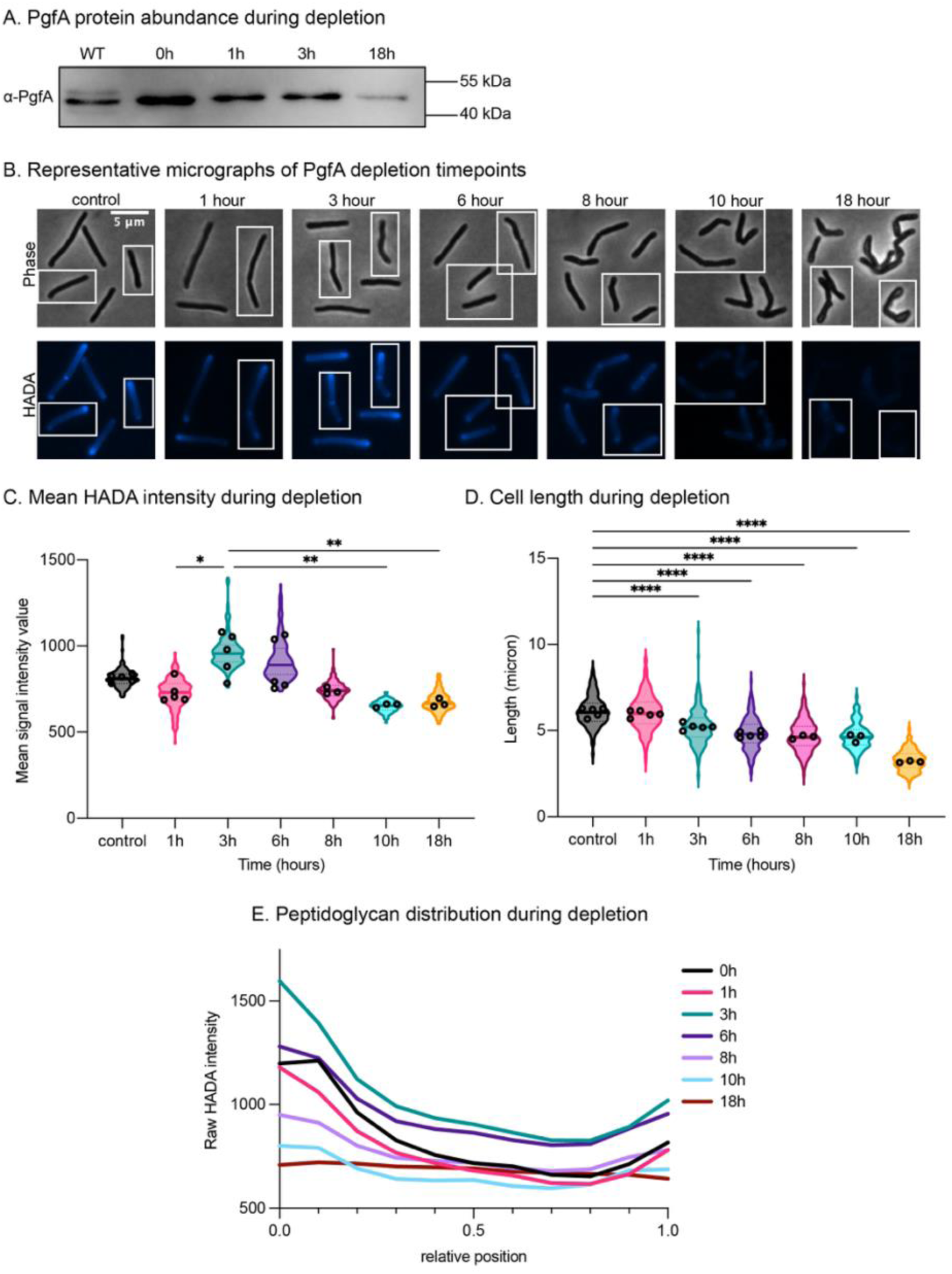
PgfA depletion reveals a role in activating PG metabolism at the poles. *(A)* Western blot showing PgfA protein levels during CRISPRi-mediated depletion. Wild-type mc^2^155 serves as control. PgfA detected at ∼45 kDa. Equal amounts of total protein were loaded per lane. Samples, in order: mc^2^155 (wild-type), 0h (depletion control), 1h, 3h, and 18h post-anhydrotetracycline (ATc) induction. *(B)* Representative phase-contrast and HADA fluorescence micrographs of cells during PgfA depletion. Cells were grown in the presence of ATc to induce CRISPRi-mediated transcriptional repression of *pgfA*, and cells were stained with HADA and imaged at indicated timepoints. Scale bar, 5 µm. *(C)* Mean HADA intensity during PgfA depletion. Violin plots show median and quartiles. Statistical comparisons are between timepoints as shown. Every dataset represents n=>300, >100 per biological replicate. (* p < 0.05, ** p < 0.01, one-way ANOVA with Tukey’s multiple comparisons). Where no line is present, p value > 0.05. *(D)* Cell length during PgfA depletion. Violin plots show median and quartiles. Statistical comparisons are to control (0h). (**** p < 0.0001, one-way ANOVA with Tukey’s multiple comparisons). *(E)* Spatial distribution of peptidoglycan metabolism during PgfA depletion. Relative HADA intensity profiles along normalized cell length (0 = old pole, 1 = new pole) at each time point. Lines represent mean intensity at each position. All data represents n=>300 cells, >100 per biological replicate. Statistical tests were performed to compare the replicate means.

Analysis of PG spatial distribution revealed a progressive decrease in polar labeling (Fig. 2E). In control cells, HADA labeling showed the characteristic polar labeling pattern, with the highest signal at the old pole. During depletion this polar enrichment pattern gradually flattened. By 18h the cells exhibited uniformly low PG labeling along their length and significantly decreased overall intensity. The eventual collapse of polar PG metabolism is consistent with loss of PgfA’s essential function in cell wall synthesis. We note that the pattern of cell morphology and HADA distribution at the 18h depletion timepoint is the same as that seen for cells depleted for Wag31(49–51) and MmpL3(45). The increase in HADA incorporation at 3h is also notable and shows that as PgfA levels decline, cells transiently experience elevated PG metabolism before eventually collapsing.

### PgfA overexpression globally dampens peptidoglycan metabolism

Our finding that PgfA depletion causes a transient increase in peptidoglycan metabolism at 3h suggests PgfA may have an inhibitory function. If PgfA can inhibit PG synthesis, we would expect PgfA overexpression to decrease PG metabolism. To test this, we assessed how increasing PgfA levels affects PG and mycolic acid (MA) metabolism.

We first determined how overexpression of PgfA impacted its distribution. When expressed under the native promoter as a single copy, PgfA-mRFP localized predominantly to cell poles as previously reported (1). When expressed as a merodiploid under a strong promoter, we observed increased fluorescence intensity throughout the cell (Fig. 3A). Using *M.smegmatis* mc^2^155 as our control and our overexpression strain containing PgfA expressed as a merodiploid under a strong promoter, we analyzed the spatial patterns of cell wall synthesis using fluorescent metabolic labeling. In wild-type cells, both PG (HADA) and MA (DMN-trehalose) incorporation were strongest at the cell poles (Fig. 3B and 3D).

**Figure 3.**
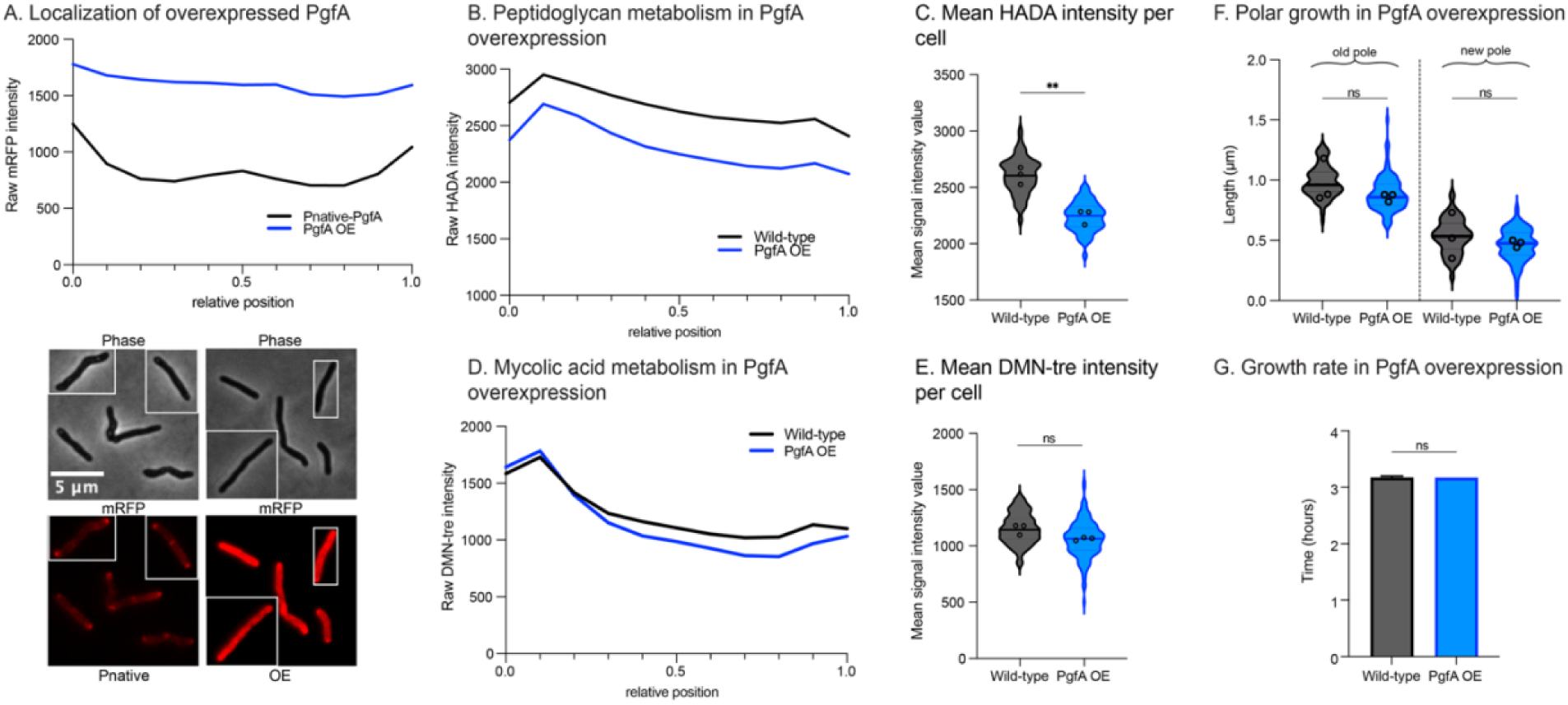
PgfA overexpression globally dampens peptidoglycan metabolism. (A) Localization of PgfA-mRFP as a single copy under its native promoter (Pnative-PgfA) vs PgfA-mRFP expressed as a merodiploid under a strong promoter causing overexpression (PgfA OE). Lines are the mean intensity along the length of >300 cells with the old pole set to 0 and the new pole set to 1 on the Y axis. Poles were assigned based on HADA intensity. **Top**: Fluorescence intensity profiles. **Bottom**: Representative phase-contrast and fluorescence images. (B) Peptidoglycan metabolism in PgfA overexpression. Fluorescence intensity profiles of HADA labeling along cell length in PgfA OE (blue) compared to wild-type (black). Graph organized as in *(A)*. All data represents n=>300 cells, >100 per biological replicate. (C) Quantification of mean HADA intensity per cell from experiment in *(B)*. (** p < 0.01, unpaired t-test). Violin plots show median and quartiles. All data represents n=>300 cells, >100 per biological replicate. (D) Mycolic acid metabolism in PgfA overexpression. Fluorescence intensity profiles of DMN-trehalose labeling along cell length in PgfA OE compared to wild-type. Graph organized as in *(A)*. All data represents n=>300 cells, >100 per biological replicate. (E) Quantification of mean DMN-trehalose intensity per cell from experiment in *(D).* (*ns*, p > 0.05, unpaired t-test). Violin plots show median and quartiles. All data represents n=>300 cells, >100 per biological replicate. (F) Quantification of polar elongation in wild-type vs PgfA overexpression. New polar growth was imaged at both poles following pulse-chase-pulse labeling and lengths were manually measured and quantified using ImageJ software. All data represents n=>300 cells, >100 per biological replicate. (*ns* p > 0.05, unpaired t-test). (G) Growth rate comparison between wild-type and PgfA overexpression strains. Doubling times were measured in 7h9 medium with no antibiotics. Bar graph shows mean ± s.d of three biological replicates. (*ns*, p > 0.05, unpaired t-test).

PgfA overexpression caused a marked reduction in overall HADA intensity (Fig. 3C), indicating decreased PG metabolism. The spatial distribution of HADA remained polar despite the overall reduction (Fig. 3B), suggesting PgfA overexpression broadly reduces PG metabolism rather than specifically affecting polar synthesis. In contrast, DMN-trehalose incorporation showed no significant change (Fig. 3E), with polar localization maintained (Fig. 3D).

To quantitatively assess polar elongation, we performed pulse-chase-pulse labeling. PgfA overexpression did not significantly impair polar elongation compared to wild-type cells (Fig. 3F). Similarly, growth rate in PgfA-overexpressed cells was not significantly different from wild-type cells (Fig. 3G). Together, these findings demonstrate that elevated PgfA levels specifically inhibit PG metabolism without affecting mycolate metabolism. HADA stains both remodeled and newly synthesized peptidoglycan (27); the change in HADA signal without a change in cell growth suggests that PgfA overexpression downregulates peptidoglycan remodeling.

### Decreased periplasmic TMM levels inhibit polar growth

Having established that PgfA impacts peptidoglycan metabolism, we sought to understand what signals regulate PG synthesis at the poles. In a previous study, we found that blocking TMM transport by treatment with 1X MIC of NITD-349 decreased PG metabolism as measured by HADA labeling (45). NITD-349 inhibits the TMM transporter MmpL3, causing a decrease in periplasmic TMMs (52).

To explore whether TMMs might have a specific role in regulating cell wall metabolism, we sought to determine whether non-inhibitory concentrations of NITD-349 could impact polar growth. As controls, we used meropenem (PG synthesis inhibitor targeting PBPs and L,D-transpeptidases)(53) and rifampin (transcription inhibitor)(54).

First, we determined the highest concentration of each drug that did not inhibit the doubling time in a standard growth curve (Fig. 4A), allowing us to assess specific effects on cell wall synthesis independently of bulk growth perturbations. Green arrows represent the noninhibitory antibiotic concentrations used for the following assay.

**Fig. 4.**
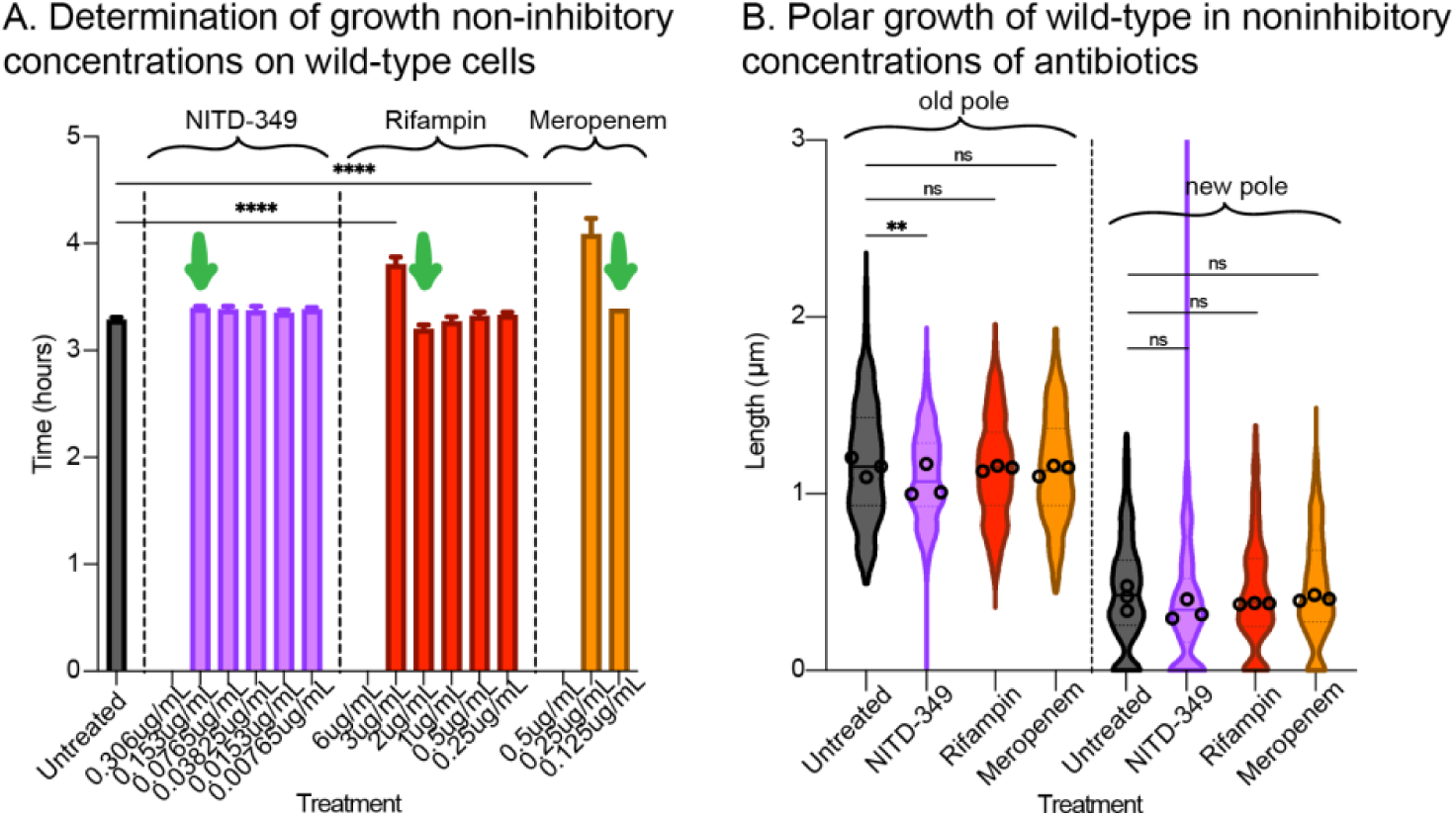
Decreased periplasmic TMM levels inhibit polar growth *(A)* Determination of growth noninhibitory concentrations of NITD-349, rifampin, and meropenem on wild-type cells. Antibiotic concentrations that inhibited growth entirely are shown without a bar. The following final concentrations of NITD-349 (0.153 µg ml⁻¹), rifampicin (2 µg ml⁻¹), or meropenem (0.125 µg ml⁻¹) were determined to be the highest which did not perturb bulk growth rate. Data represents the mean ± s.d. of three biological replicates. (*ns* p > 0.05, **** p < 0.0001, one-way ANOVA with Tukey’s multiple comparisons). *(B)* Quantification of polar elongation in wild-type under the same treatment conditions as *(A)*. Poles were assigned based on HADA intensity – the old pole is assumed to be the pole that is brighter. Violin plots show individual cell distributions with mean (dashed line) and interquartile range (dotted line). All microscopy data represents n=>300 cells, >100 per biological replicate. (*ns* p > 0.05, ** p < 0.01, one-way ANOVA with Tukey’s multiple comparisons).

To quantitatively assess polar elongation, we treated Msmeg cells with non-inhibitory concentrations of each drug and performed pulse-chase-pulse labeling with sequential Fluorescent D-amino acids (FDAAs) HADA and NADA(55). Cells were stained with HADA first, outgrown in fresh media with antibiotics, then stained with NADA and imaged immediately. Polar elongation was measured by quantifying the length of the HADA-free region at the poles. Despite unchanged bulk growth rates, NITD-349 treatment reduced polar elongation at the old pole, whereas rifampin and meropenem showed no effect (Fig. 4B). This selective impairment of polar growth suggests that periplasmic TMM availability has a specific role in regulating polar growth.

### PgfA overexpression and NITD-349 treatment show epistatic effects on peptidoglycan metabolism

Our findings demonstrate that both PgfA overexpression (Fig. 3) and NITD-349 treatment (45) reduce peptidoglycan metabolism, specifically at the old pole (Fig. 4B). Because PgfA binds TMMs(1) and NITD-349 blocks TMM transport(52), we hypothesized that both perturbations inhibit PG metabolism by altering the availability of TMM to PgfA. Specifically, we reasoned that overexpression of PgfA without a corresponding increase in TMM flux may drive PgfA into a TMM-free state that inhibits PG metabolism. To test this model, we treated PgfA-overexpressing cells with 1X MIC NITD-349 and again measured PG synthesis by HADA incorporation. If PG inhibition results from PgfA existing in a TMM-free inhibitory state, then further depletion of TMM by NITD-349 should not further reduce PG metabolism in cells overexpressing PgfA. As controls to test pathway specificity, we included ebselen,which blocks downstream mycolate incorporation, (56) and meropenem, which inhibits PG crosslinking. Consistent with our hypothesis, PgfA overexpression in NITD-349-and ebselen-treated cells showed no additional reduction in HADA incorporation compared to the drug treatments alone (Fig. 5A). Similarly, ebselen-and NITD-349 drug treatments do not further decrease HADA labeling in the PgfA overexpression background (Fig. 5A). These data suggest that ebselen and NITD-349 impact PG metabolism through the same pathway as PgfA overexpression. In contrast, meropenem treatment showed near-additive effects with PgfA overexpression, with dual treatment causing a significantly greater reduction in PG labeling than either treatment alone (Fig. 5A). This result indicates that PgfA and meropenem act through independent mechanisms, as expected since meropenem directly inhibits transpeptidase activity.

**Figure 5.**
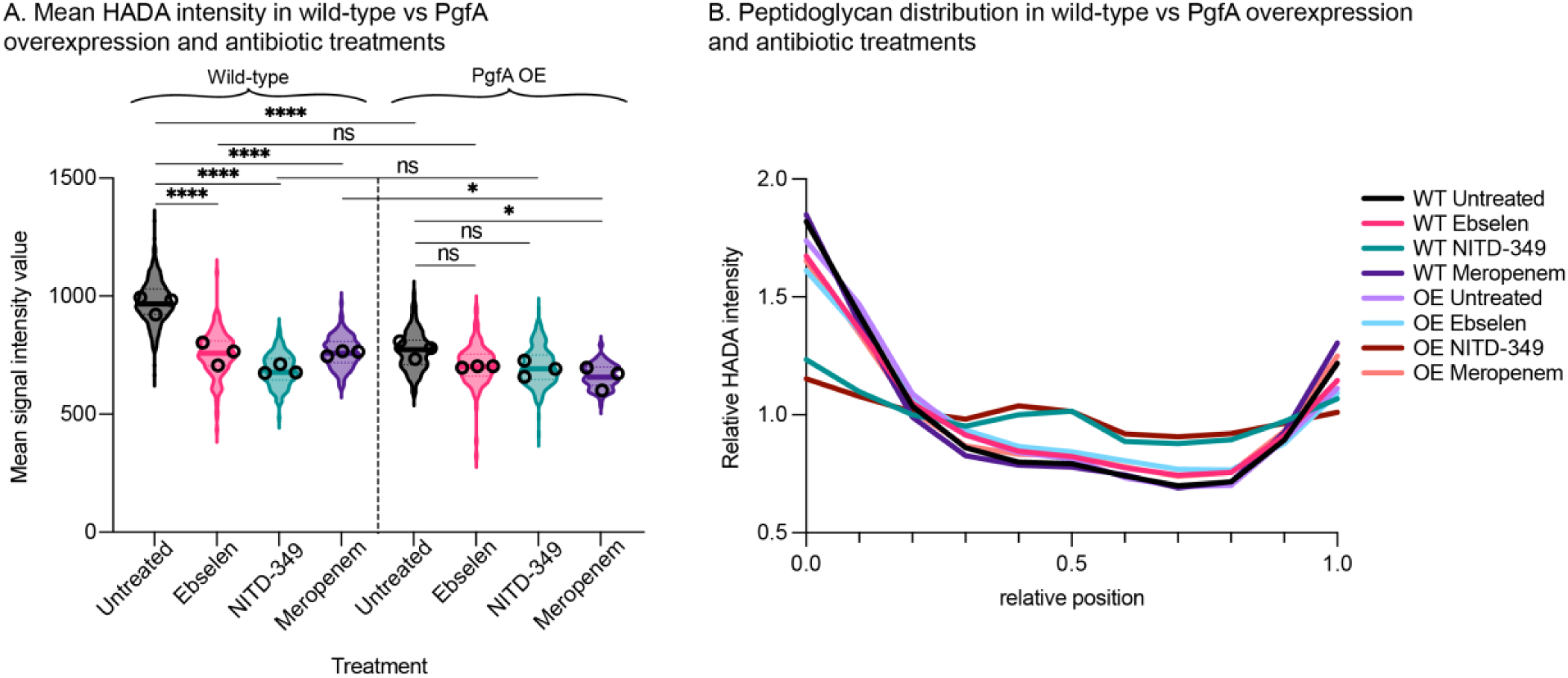
PgfA overexpression and NITD-349 treatment show epistatic effects on peptidoglycan metabolism. *(A)* Mean HADA fluorescence intensity in wild-type *M. smegmatis* cells vs PgfA overexpressed cells treated with 1X MIC of NITD-349 (1.53 µg ml⁻¹), ebselen (20 µg ml⁻¹), and meropenem (2.5 µg ml⁻¹) for 1 hour. All data represents the mean ± s.d. of three biological replicates, and small black circles represent the means per biological replicate. (* p < 0.05, **** p < 0.0001, one-way ANOVA with Tukey’s multiple comparisons). All data represents n=>300 cells, >100 per biological replicate. Statistical tests were calculated based on the replicate means. *(B)* Spatial distribution of peptidoglycan metabolism for wild-type and PgfA overexpression cells treated as in *(A)*. Relative fluorescence intensity profiles show HADA signal normalized to the mean intensity of each cell. All data represents n=>300 cells, >100 per biological replicate.

Notably, NITD-349 and ebselen produced strikingly different effects on the spatial organization of PG metabolism despite both reducing overall HADA intensity. NITD-349 caused complete loss of polar PG labeling in both wild-type and PgfA OE backgrounds. Meanwhile, ebselen reduced overall PG metabolism but preserved polar organization regardless of PgfA levels (Fig. 5B, Supplemental Fig. 2A).

Overall, these results suggest that changes in periplasmic TMM levels affect peptidoglycan metabolism through the same pathway as PgfA. Ebselen and NITD-349 both cause a decrease in PG metabolism; however, only NITD-349 – which should decrease periplasmic TMM levels – downregulates polar peptidoglycan metabolism. Ebselen may briefly increase periplasmic TMM levels, before it causes destruction of the outer membrane. These data support the idea that periplasmic TMMs stimulate polar peptidoglycan metabolism, and are connected to the function of PgfA.

## Discussion

Our findings establish that both PgfA and periplasmic TMM levels affect peptidoglycan metabolism. Changes in PgfA levels reveal that PgfA can have both inhibitory and stimulatory effects on PG metabolism (Fig. 2,3), though our data do not indicate whether this is direct or indirect. The epistasis between PgfA overexpression and NITD-349 treatment (Fig. 5), provides strong evidence that these perturbations work through a shared pathway.

We propose a model where TMMs regulate PG metabolism through their interaction with PgfA (1). We hypothesize that TMM-bound PgfA functions as an activator of PG metabolism and polar growth, while TMM-free PgfA inhibits PG metabolism (Fig. 6).

**Figure 6.**
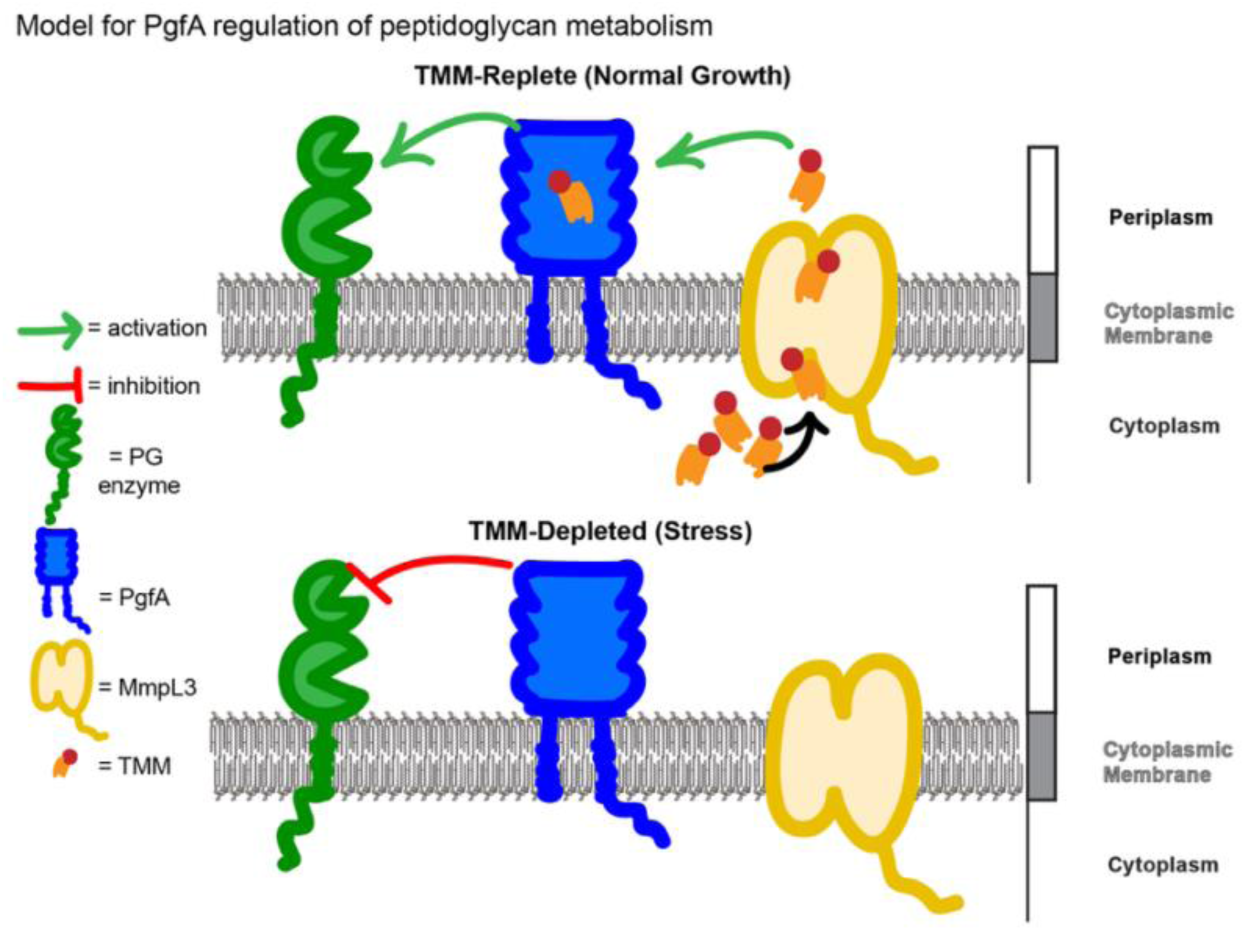
Model for PgfA regulation of peptidoglycan Model for PgfA regulation of peptidoglycan metabolism. **Top:** In wild-type cells, PgfA localizes to cell poles where MmpL3 delivers TMM. PgfA protein is TMM-bound, allowing normal polar growth. **Bottom:** TMM-free PgfA inhibits peptidoglycan metabolism. Green arrows show activation, red bars show inhibition.

PgfA overexpression likely increases the pool of TMM-free PgfA that inhibits PG metabolism while spatial organization is maintained (Fig. 3) because Pks13 (13) and MmpL3 (11) continue delivering TMM to the poles. In this model, NITD-349 treatment creates abundant TMM-free PgfA by blocking TMM transport, causing both PG inhibition and loss of polarity due to disrupted polar TMM delivery, consistent with the epistatic experiments and HADA labeling shown in Figure 5.

We find that PgfA overexpression does not significantly affect polar elongation at either pole in wild-type cells. In contrast, PgfA, but not MmpL3 over-expression, increases elongation at the old pole in Δ*lamA* cells (1). As LamA is required for PgfA and MmpL3 polar localization, overexpressing PgfA in Δ*lamA* cells likely forces a fraction of the protein to the poles (Fig. 3A), even in the absence of LamA-mediated recruitment. That PgfA is able to restore growth in Δ*lamA* further suggests that PgfA is a critical driver of cell wall expansion and elongation at the growing pole.

To our knowledge, PgfA represents the first identified regulatory protein that could coordinate PG and MA synthesis by sensing the availability of one pathway product to regulate another pathway in mycobacteria. Previous work established that the scaffolding protein Wag31 organizes cell wall synthesis machinery at the cell poles (13, 45), and that the distribution of this machinery at the two poles is partly regulated by LamA (1, 57). Our recent work suggests that the recruitment and regulation of MmpL3 by Wag31 is a key upstream event that regulates cell wall metabolism at both poles (45). The importance of TMM transporter MmpL3 in regulating polar growth can be partly explained with our model that TMMs are an upstream signal that activates metabolism of the inner layer of the cell wall, likely through PgfA. While our observation that PgfA also has a role inhibiting PG metabolism is new, previous work showed that PgfA overexpression inhibits the production of lipoarabinomannans (LAM) and lipomannans (LM) (1). It is entirely likely that these inhibitory functions are linked; perhaps there is a system to coordinate LM and LAM with PG synthesis.

Cell wall precursors are detected by various systems to help regulate cell wall metabolism across bacterial clades. In *Staphylococcus aureus* and *M. tuberculosis*, cell wall regulatory kinase PknB is activated by binding to periplasmic lipid II through its extracytoplasmic domain (58–60). In Gram-positive *B. subtilis*, the availability of lipid II is a major regulator of cell elongation machinery (61, 62). In Gram-negative *Pseudomonas aeruginosa*, MraY is allosterically inhibited by extracytoplasmic lipid II(63), and PG precursor enzyme MurA is inhibited by its substrate (64) and also helps to coordinate LPS synthesis (65). These reports demonstrate bacteria possess diverse mechanisms to sense precursor availability to balance envelope synthesis. We believe PgfA may represent a regulatory mechanism similar to those described above, one where its TMM binding controls its functional state. If correct, this would allow reversible, metabolite-responsive regulation that could rapidly adapt to changing conditions. Further biochemical and structural studies will be required to establish the molecular mechanism by which PgfA coordinates these pathways, particularly to identify which PG synthases PgfA regulates and how conformational changes of its periplasmic domain drive the functional switch.

Understanding these coordination mechanisms has implications for antibiotic development: many antimycobacterial drugs target individual envelope pathways (β-lactams, isoniazid, ethambutol), but coordination proteins like PgfA may enable compensatory responses to drug treatments. Targeting checkpoint proteins could enhance existing antibiotics or reveal combination therapy strategies.

## Materials and Methods

### Bacterial strains and growth conditions

All *M.smegmatis* strains were grown in 7H9 (BD, Sparks, MD) liquid medium with 0.2% glycerol, 0.05% Tween80, and ADC (5 g/L albumin, 2 g/L dextrose, 0.85 g/L NaCl, and 0.003 g/L catalase) added. *M.smegmatis* strains were also grown on LB Lennox plates. TOP10 *E.coli* competent cells were used for gene cloning. Antibiotic concentrations for *M.smegmatis* were as follows: hygromycin – 50 μg/mL; nourseothricin – 20 μg/mL; kanamycin – 25 μg/mL; zeocin – 20 μg/mL; streptomycin – 20 μg/mL. Antibiotic concentrations for *E.coli* were as follows: nourseothricin – 40 μg/mL; kanamycin – 50 μg/mL; zeocin – 25 μg/mL.

### Strain construction

Strains used in this study are listed in Table S1. The PgfA allele swap strain, PgfA overexpression strain, and native PgfA localization strain were provided to us by the Rego lab, as described(1). The PgfA and MmpL3 depletion strains were made using *E.coli* CRISPRi plasmids obtained from the Mycobacterial Systems Resource Database(29). All other plasmids were made using Gibson cloning (66) and transformed into either wild-type *M. smegmatis* or the PgfA allele swap strain. Primers are listed in Table S2.

### Drug treatment assays

All microscopy with drug treatments were performed using log phase cells in 7H9 media at the following concentrations: rifampin 2 µg/mL; meropenem 2.5 µg/mL (for HADA microscopy) or 0.125 µg/mL (for noninhibitory polar elongation); NITD-349 1.53 μg/mL (for localization and HADA microscopy) or 0.153 μg/mL (for noninhibitory polar elongation); ebselen 20 µg/mL. For polar elongation microscopy experiments, untreated cells were incubated at 37°C while rolling for 45 mins in noninhibitory concentrations of the respective drugs, then HADA was added for the remaining 15 minutes. Cells were washed with 7H9 once, then immediately mounted on 1.5% HdB complete agarose pads and imaged (for noninhibitory polar elongation). For HADA microscopy experiments, untreated cells were incubated at 37°C while rolling for 45 mins in 1X MIC concentrations of the respective drugs, then HADA was added for the remaining 15 minutes. Cells were washed once with 1X PBS, fixed for 10 minutes at 25°C with 1.6% paraformaldehyde in 1X PBS, then washed once more with 1X PBS, mounted on 1.5% PBS agarose pads and imaged.

### Growth Curves

Cultures which had been in logarithmic phase for at least 24 hours were diluted to OD_600_ = 0.1 in 96-well plates. No antibiotics were used during growth curves. A Synergy Neo2 linear multi-Mode Reader was used to shake the plates at 37°C, and the OD_600_ was read every 15 min.

### Cell staining

For all microscopy involving cell wall labeling, cells were in logarithmic phase (OD 0.4) prior to incubation with the fluorescent D-amino acid dye. For HADA(55) (BioTechne Cat #6647) and NADA(67) (BioTechne Cat #6648), cells were incubated for 15 minutes at 37°C while rolling at a final concentration of 10 µM each, then the dye was washed out by centrifugation and resuspension in plain media. For DMN-trehalose(32), cells were incubated for 15 minutes at 37°C while rolling at a final concentration of 1 µg/mL. After washing out the dye, cells were resuspended in either 7H9, HdB with and without Tween80 or glycerol, or 1X PBS with tyloxapol. All cells were imaged live.

### Elongation assays

1mL of log. phase cells (OD 0.4) were stained with HADA, then washed once with 7H9. The pellet was resuspended in 1mL of fresh 7H9 and outgrown for 45 minutes at 37°C while rolling. The cells were then stained with NADA at a final concentration of 10 µM for 15 minutes at 37°C while rolling, washed once more with 7H9, mounted on 1.5% HdB complete agarose pads, and imaged.

### Starvation assays

Log. phase cells grown in 7H9 were pelleted and washed once with either HdB with 0.05% tween80 and no glycerol (for soft starvation) or PBS with 0.05% tyloxapol (for hard starvation) before being resuspended in their respective media and incubated at 37°C while rolling for either 1, 3, or 16 hours. Control samples were grown in 7H9, kept in logarithmic phase, and imaged at the respective times alongside starved cells.

### CRISPRi depletion assays

Transcriptional repression was induced in log. phase cells (OD 0.4) using anhydrotetracycline (ATc) at a concentration of 500ng/mL. Samples from a bulk culture were collected and stained with HADA at 0h (control), 1h, 3h, 6h, 8h, 10h, and 18h. At each timepoint, cells were stained with HADA as described above and imaged live.

### Microscopy

All microscopy was performed on live cells mounted on 1.5% Hartmans-de Bont (HdB) agarose pads (for fed or soft starvation experiments) or 1.5% PBS agarose pads (for hard starvation experiments). Agarose pads made with HdB media containing both 0.05% Tween80 and glycerol (HdB complete) were used for all non-starvation experiments.

Agarose pads made with HdB media with only 0.05% Tween80 and no glycerol were used for imaging soft starved cells. Agarose pads made with 1X PBS containing 0.05% tyloxapol were used for imaging hard starved cells. Cells were imaged using a Nikon Ti-2 widefield fluorescence microscope with a Plan Apo 100×, 1.45 NA objective. A Photometrics Prime 95B camera was used to collect images. NADA was imaged using a filter cube with a 470/40 nm bandpass excitation filter, a 495 nm dichroic mirror, and a 525/50 nm emission filter. mRFP was imaged using a filter cube with a 560/40 nm bandpass excitation filter, a 585 nm dichroic mirror, and a 630/70 nm emission filter. HADA was imaged using a filter cube with a 350/50 nm bandpass excitation filter, a 400 nm dichroic mirror, and a 460/50 nm emission filter.

### Microscopy analysis

Cell segmentation and fluorescence quantification were performed using MicrobeJ(68) (version 5.13I) in ImageJ/FIJI. For fluorescence intensity profiles, cell length was normalized (0 = old pole, 1 = new pole) and fluorescence intensity was measured and averaged at 10 positions along the long axis of the cell. For polar elongation measurements, the old pole was assumed to be the one with the brighter HADA signal. The lengths of new pole growth, identified by NADA-labeled regions, were measured manually for each cell. Width range was calculated as the difference between maximum and minimum width for each cell.

Data were exported from MicrobeJ and graphed using GraphPad Prism version 10.6.1.

## Statistical analysis

All experiments were performed with three biological replicates. For microscopy experiments, at least 100 cells per replicate were analyzed (n≥300 total cells per condition). For growth curve experiments, three technical replicates were performed for each biological replicate. Statistical comparisons were performed as indicated in figure legends using GraphPad Prism version 10.6.1. P-values: ns, p > 0.05; *, p < 0.05; **, p < 0.01; ***, p < 0.001; ****, p < 0.0001. All data are presented as violin plots with median (solid line) and quartiles (dotted lines), or as bar graphs with mean ± standard deviation. Correlation between PgfA-mRFP and HADA fluorescence intensities was assessed using Spearman rank correlation. For polar analysis, mean intensities were extracted from old pole (normalized positions 0-0.025) and new pole (0.975-1.0) regions using previously published code(69) that was slightly altered. For midcell analysis, all intensity measurements within the midcell region (0.4-0.6) were used. Statistical analyses were performed using MATLAB and Python. Bespoke Python codes for the midcell enrichment analysis and generation of rank correlation values for either the poles or the midcell are available in Supplementary Figure 3 and Supplementary Figures 4 and 5 respectively.

## Acknowledgements

We thank Neda Habibi Arejan for the max pole intensity analysis MATLAB code. We thank Ei Phoo Phoo Aung, Keith Derbyshire and Joe Wade for the generous gift of the MSRdb collection.

This work was funded by grant R01AI148917 to CCB, 1R01AI148255 to EHR, and 1R01AI148917-S1 to KET.

