## Supplement for "Polar growth factor PgfA regulates polar peptidoglycan synthesis as well as mycolate synthesis in *Mycobacterium smegmatis*"

**Supplemental Table 1. Strain list.**

| Strain number | Genotype | Source | Figure panel |
|---|---|---|---|
| 1278 | Wild-type mc2155 | | Figure 4, 3B-G, 5 |
| 3433 | $mc^2155$ L5::Pnative-Msm PgfA-mRFP-myc-nuoR ΔpgfA::zeoR | Rego Lab(1) | Figure 1, 3A |
| 3438 | $mc^2155$/ L5::pTetO-Msm PgfA-myc | Rego Lab(1) | Figure 3B-G, 5 |
| 3525 | $mc^2155$ L5::pTetO-WT PgfA-mRFP-myc-nuoR | | Figure 3A |
| 3816 | $mc^2155$/ L5::pMSMEG_0317 (pgfA).sgRNA | | Figure 2 |

**Supplemental Table 2. Plasmid list.**

| Strain number | Genotype | Source | Used in strain | Figure panel |
|---|---|---|---|---|
| 3439 | Dh5α/ pL5-pTetO-Msm PgfA-myc-nuo | Rego Lab(1) | 3438 | Figure 3B-G, 5 |
| 3806 | Dh5α/ pMSMEG_0317 (pgfA).sgRNA) | MSRdb(2) | 3816 | Figure 2 |
| 3506 | Top10/ pL5-pTetO-Msm WT PgfA-mRFP-myc | | 3525 | Figure 3A |

**Supplemental Table 3. Primer list.**

| Strain number | Feature | Primer sequence |
|---|---|---|
| 3506 | Top10/ pL5-pTetO-Msm WT PgfA-mRFP-myc | Fwd:<br>AGATTCGCCGCCCGAAATGAGCACGATCCGCATGCTTAATTAAGAAGGAGATATACTAttgAACCGCGCTGTGGCGCTGCGT<br>Add mRFP to FL PgfA:<br>CACCGGACCGACCGATCggcggctcgGCCTCCTCCGAGGAC<br>Add mRFP to FL PgfA:<br>ttgatgaCGTCCTCGGAGGAGGCcgagccgccGATCGGTCGGTCCGG<br>Rev:<br>GGTCCCCAATTAATTAGCTAAAGCTTTCACAGGTCTTCCTCGCTGATCAGCTTCTGCTCGGCGCCGGTGGAGTGgcggccctcggcgc |

### Supplemental Figures

### Supplemental Figure 1. PgfA-mRFP localization in log phase and complete carbon starvation

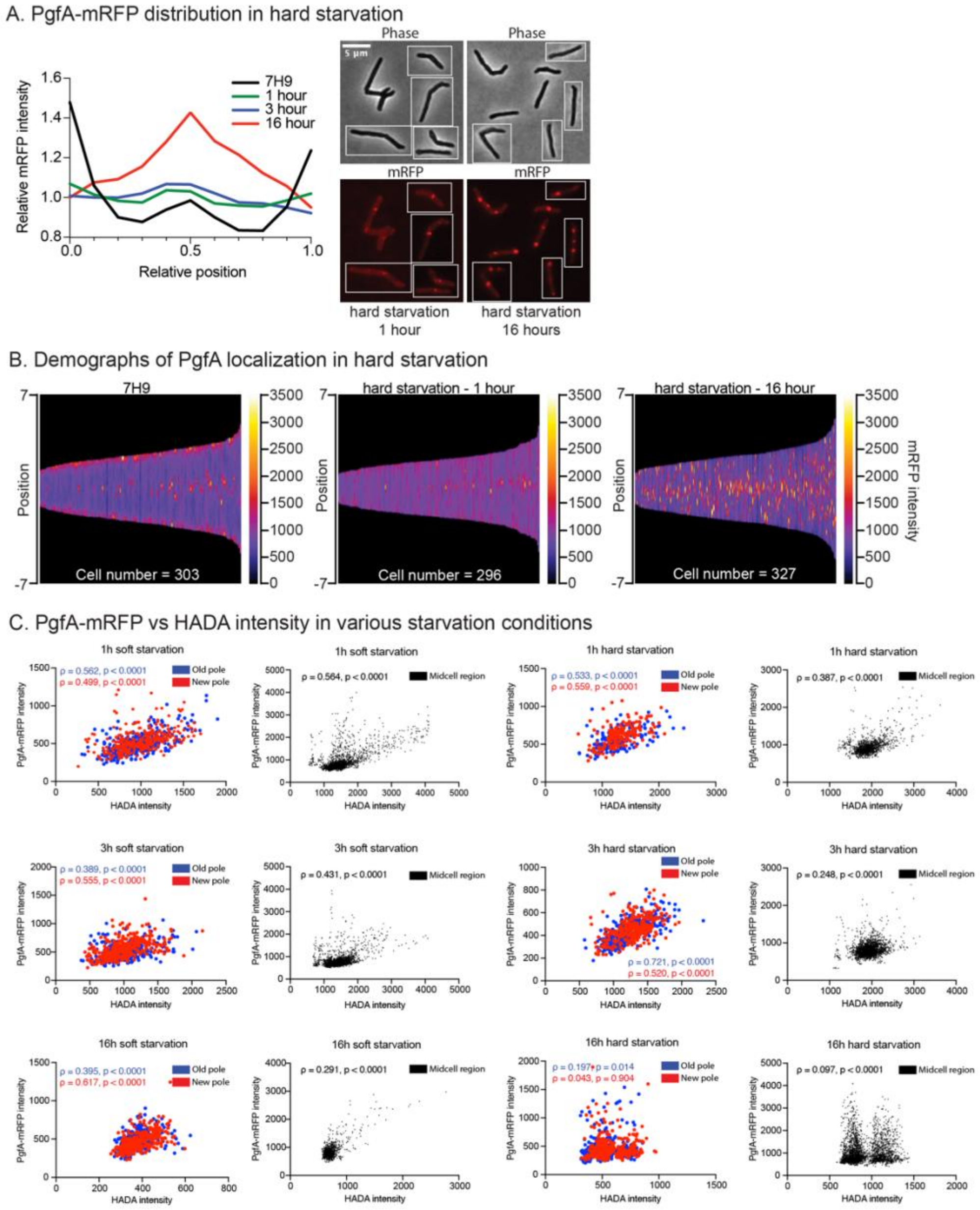

**Supplemental Figure 2. Raw peptidoglycan distribution in wild-type vs PgfA overexpression and antibiotic treatments**

A. Raw peptidoglycan distribution in wild-type vs PgfA overexpression and antibiotic treatments

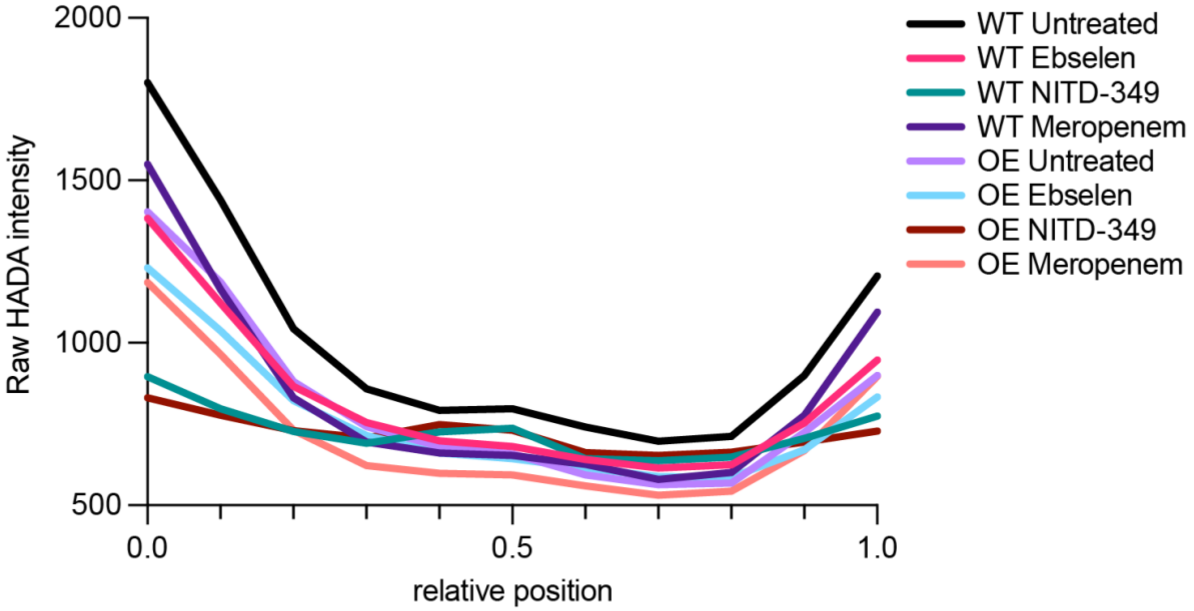

**Supplemental Figure 3. Septal PgfA-mRFP and HADA colocalization analysis python code.**

```
#!/usr/bin/env python3
"""
Septal Colocalization Analysis for GraphPad Prism
Analyzes mRFP and HADA colocalization at cell septa (midcell, position
0.4-0.6)
"""

import pandas as pd
import numpy as np
from scipy import stats

#
======================================================================
======
# CONFIGURATION
#
======================================================================
======
INPUT_FILE = 'data.xlsx'
SEPTAL_START = 0.4  # Start of septal region (normalized position)
SEPTAL_END = 0.6    # End of septal region (normalized position)
PEAK_DISTANCE_THRESHOLD = 0.1  # Consider peaks colocalized if within
this distance

#
======================================================================
======
# READ DATA
#
======================================================================
======
print("=" * 80)
print("SEPTAL COLOCALIZATION ANALYSIS")
print("=" * 80)

df = pd.read_excel(INPUT_FILE)

# Clean up column names. Change '_____' to your gene name & edit for
your cell
# Make sure to replicate changes throughout
df = df.rename(columns={
    'CELL ID': 'Cell_ID_HADA',
    'X - HADA': 'Position_HADA',
    'Y - HADA': 'HADA_Intensity',
    'CELL ID.1': 'Cell_ID_mRFP',
    'X - ____': 'Position_mRFP',
    'Y - ____': 'mRFP_Intensity'
})

# Use HADA position and cell ID (they should be identical)
```

```
df['Position'] = df['Position_HADA']
df['Cell_ID'] = df['Cell_ID_HADA']

print(f"\nTotal datapoints: {len(df)}")
print(f"Unique cells: {df['Cell_ID'].nunique()}")

#
======================================================================
======
# FILTER FOR SEPTAL REGION (0.4 - 0.6)
#
======================================================================
======
septal_df = df[(df['Position'] >= SEPTAL_START) & (df['Position'] <=
SEPTAL_END)].copy()

print(f"\nSeptal datapoints (positions {SEPTAL_START}-{SEPTAL_END}):
{len(septal_df)}")
print(f"Cells with septal data: {septal_df['Cell_ID'].nunique()}")

#
======================================================================
======
# CALCULATE HADA THRESHOLD
#
======================================================================
======
# Define HADA-positive as mean + 1 SD
hada_mean = septal_df['HADA_Intensity'].mean()
hada_std = septal_df['HADA_Intensity'].std()
hada_threshold = hada_mean + hada_std

septal_df['HADA_above_threshold'] = septal_df['HADA_Intensity'] >
hada_threshold

print(f"\nHADA threshold: {hada_threshold:.2f} (mean + 1 SD)")
print(f"Datapoints above threshold:
{septal_df['HADA_above_threshold'].sum()}")

#
======================================================================
======
# PER-CELL ANALYSIS
#
======================================================================
======
print("\n" + "=" * 80)
print("CALCULATING PER-CELL METRICS")
print("=" * 80)
```

```
cell_metrics_list = []

for cell_id, cell_data in septal_df.groupby('Cell_ID'):
    if len(cell_data) < 2:  # Need at least 2 points for correlation
        continue

    # Pearson correlation
    pearson_r =
cell_data['HADA_Intensity'].corr(cell_data['mRFP_Intensity'])
    r_squared = pearson_r ** 2 if not pd.isna(pearson_r) else 0

    # Find peak positions
    hada_peak_idx = cell_data['HADA_Intensity'].idxmax()
    mrfp_peak_idx = cell_data['mRFP_Intensity'].idxmax()

    hada_peak_pos = cell_data.loc[hada_peak_idx, 'Position']
    mrfp_peak_pos = cell_data.loc[mrfp_peak_idx, 'Position']

    # Peak distance
    peak_distance = abs(hada_peak_pos - mrfp_peak_pos)
    peaks_colocalized = peak_distance < PEAK_DISTANCE_THRESHOLD

    # mRFP enrichment at HADA-positive positions
    hada_positive = cell_data[cell_data['HADA_above_threshold']]
    hada_negative = cell_data[~cell_data['HADA_above_threshold']]

    if len(hada_positive) > 0 and len(hada_negative) > 0:
        mrfp_at_hada_peaks = hada_positive['mRFP_Intensity'].mean()
        mrfp_at_hada_neg = hada_negative['mRFP_Intensity'].mean()
        enrichment = mrfp_at_hada_peaks / mrfp_at_hada_neg if
mrfp_at_hada_neg > 0 else np.nan
    else:
        enrichment = np.nan

    cell_metrics_list.append({
        'Cell_ID': cell_id,
        'Pearson_r': pearson_r,
        'R_squared': r_squared,
        'HADA_peak_position': hada_peak_pos,
        'mRFP_peak_position': mrfp_peak_pos,
        'Peak_distance': peak_distance,
        'Peaks_colocalized': peaks_colocalized,
        'mRFP_enrichment_at_HADA_peaks': enrichment,
        'N_datapoints_in_septum': len(cell_data)
    })

cell_metrics_df = pd.DataFrame(cell_metrics_list)

#
======================================================================
```

```
======
# GLOBAL CORRELATION (ALL SEPTAL POINTS COMBINED)
#
======================================================================
======
global_pearson_r =
septal_df['HADA_Intensity'].corr(septal_df['mRFP_Intensity'])
global_r_squared = global_pearson_r ** 2

# Linear regression for plotting
slope, intercept, r_value, p_value, std_err = stats.linregress(
    septal_df['HADA_Intensity'],
    septal_df['mRFP_Intensity']
)

print(f"\nGlobal correlation (all septal datapoints):")
print(f"  Pearson r = {global_pearson_r:.3f}")
print(f"  R² = {global_r_squared:.3f}")
print(f"  p-value = {p_value:.2e}")
print(f"  Linear fit: y = {slope:.3f}x + {intercept:.2f}")

#
======================================================================
======
# SUMMARY STATISTICS
#
======================================================================
======
print("\n" + "=" * 80)
print("SUMMARY STATISTICS")
print("=" * 80)

# Remove NaN values for statistics
valid_r2 = cell_metrics_df['R_squared'].dropna()
valid_enrichment =
cell_metrics_df['mRFP_enrichment_at_HADA_peaks'].dropna()

print(f"\nPer-cell R² distribution:")
print(f"  Mean R² = {valid_r2.mean():.3f} ± {valid_r2.std():.3f}
(SD)")
print(f"  Median R² = {valid_r2.median():.3f}")
print(f"  Range: {valid_r2.min():.3f} to {valid_r2.max():.3f}")
print(f"  N cells = {len(valid_r2)}")

colocalized_count = cell_metrics_df['Peaks_colocalized'].sum()
total_cells = len(cell_metrics_df)
colocalized_pct = 100 * colocalized_count / total_cells

print(f"\nPeak colocalization:")
print(f"  Cells with colocalized peaks: {colocalized_count}/
```

```
{total_cells} ({colocalized_pct:.1f}%)")
print(f"  Mean peak distance:
{cell_metrics_df['Peak_distance'].mean():.3f} ±
{cell_metrics_df['Peak_distance'].std():.3f}")

if len(valid_enrichment) > 0:
    print(f"\nmRFP enrichment at HADA-positive regions:")
    print(f"  Mean enrichment: {valid_enrichment.mean():.2f}-fold")
    print(f"  Median enrichment: {valid_enrichment.median():.2f}-
fold")

#
======================================================================
======
# EXPORT FOR GRAPHPAD PRISM
#
======================================================================
======
print("\n" + "=" * 80)
print("EXPORTING DATA FOR PRISM")
print("=" * 80)

# 1. All septal datapoints for scatter plot
output1 = septal_df[['Cell_ID', 'Position', 'HADA_Intensity',
'mRFP_Intensity', 'HADA_above_threshold']].copy()
output1.to_csv('prism_septal_all_points.csv', index=False)
print("\n1. prism_septal_all_points.csv")
print("   → Use for scatter plot (HADA vs mRFP)")
print("   → X: HADA_Intensity, Y: mRFP_Intensity")

# 2. Per-cell metrics
cell_metrics_df.to_csv('prism_per_cell_metrics.csv', index=False)
print("\n2. prism_per_cell_metrics.csv")
print("   → Use for R² distribution, peak colocalization stats")
print("   → Plot R_squared as violin/column plot")

# 3. Summary statistics table
summary_stats = pd.DataFrame({
    'Metric': [
        'Global R²',
        'Global Pearson r',
        'Mean per-cell R²',
        'SD per-cell R²',
        '% cells with colocalized peaks',
        'Mean peak distance',
        'N cells analyzed'
    ],
    'Value': [
        f"{global_r_squared:.3f}",
        f"{global_pearson_r:.3f}",
```

```
        f"{valid_r2.mean():.3f}",
        f"{valid_r2.std():.3f}",
        f"{colocalized_pct:.1f}%",
        f"{cell_metrics_df['Peak_distance'].mean():.3f}",
        f"{len(cell_metrics_df)}"
    ]
})
summary_stats.to_csv('prism_summary_stats.csv', index=False)
print("\n3. prism_summary_stats.csv")
print("   → Summary table for your paper")

# 4. Regression data for Prism
regression_data = pd.DataFrame({
    'HADA_Intensity': septal_df['HADA_Intensity'],
    'mRFP_Intensity': septal_df['mRFP_Intensity'],
    'Fitted_line': slope * septal_df['HADA_Intensity'] + intercept
})
regression_data.to_csv('prism_regression_data.csv', index=False)
print("\n4. prism_regression_data.csv")
print("   → Use to plot regression line on scatter plot")

print("\n" + "=" * 80)
print("ANALYSIS COMPLETE!")
print("=" * 80)
print("\nTo use in GraphPad Prism:")
print("1. Import 'prism_septal_all_points.csv' for XY scatter plot")
print("2. Analyze → XY analysis → Linear regression (to get R²)")
print("3. Import 'prism_per_cell_metrics.csv' for per-cell
statistics")
print("4. Use 'prism_summary_stats.csv' for your methods/results")
```

**Supplemental Figure 4. Batch polar correlation calculator python code.**

```
#!/usr/bin/env python3
"""
Calculates Spearman correlations for old pole and new pole separately

INPUT: One Excel file with multiple sheets (one sheet per condition)
Each sheet should have columns:
  - Pole 1 P = old pole pgfA/mRFP intensity
  - Pole 1 H = old pole HADA intensity
  - Pole 2 P = new pole pgfA/mRFP intensity
  - Pole 2 H = new pole HADA intensity

OUTPUT: CSV file with correlation statistics for each condition (old
and new poles separate)
"""

import pandas as pd
import numpy as np
from scipy import stats

#
======================================================================
======
# CONFIGURATION - EDIT THIS
#
======================================================================
======

INPUT_FILE = 'pole_intensity_data.xlsx'  # <- Your Excel file name

# Column names in your Excel sheets. Change as necessary for your
gene/condition.
# Make sure to change column names throughout the rest of the code!
OLD_POLE_MRFP = 'Pole 1 P'
OLD_POLE_HADA = 'Pole 1 H'
NEW_POLE_MRFP = 'Pole 2 P'
NEW_POLE_HADA = 'Pole 2 H'

#
======================================================================
======
# EDIT CAREFULLY BELOW THIS LINE
#
======================================================================
======

print("=" * 80)
print("BATCH POLAR CORRELATION ANALYSIS")
print("=" * 80)
print(f"\nInput file: {INPUT_FILE}")
```

```
# Read Excel file
excel_file = pd.ExcelFile(INPUT_FILE)
sheet_names = excel_file.sheet_names

print(f"Found {len(sheet_names)} sheets: {sheet_names}")
print()

results = []

for sheet_name in sheet_names:
    print(f"Processing: {sheet_name}")

    # Read the sheet
    df = pd.read_excel(excel_file, sheet_name=sheet_name)

    print(f"   Rows: {len(df)}")

    #
======================================================================
==
    # OLD POLE (Pole 1)
    #
======================================================================
==

    old_pole_mrfp = df[OLD_POLE_MRFP].values
    old_pole_hada = df[OLD_POLE_HADA].values

    # Remove NaN
    mask_old = ~(np.isnan(old_pole_mrfp) | np.isnan(old_pole_hada))
    old_pole_mrfp = old_pole_mrfp[mask_old]
    old_pole_hada = old_pole_hada[mask_old]

    if len(old_pole_mrfp) >= 3:
        # Pearson
        pearson_r_old, pearson_p_old = stats.pearsonr(old_pole_hada,
old_pole_mrfp)

        # Spearman
        spearman_r_old, spearman_p_old =
stats.spearmanr(old_pole_hada, old_pole_mrfp)

        # Kendall
        kendall_tau_old, kendall_p_old =
stats.kendalltau(old_pole_hada, old_pole_mrfp)

        print(f"   📊 OLD POLE:")
        print(f"      n = {len(old_pole_mrfp)}")
        print(f"      Pearson:  r = {pearson_r_old:.4f}, r² =
```

```
{pearson_r_old**2:.4f}")
        print(f"      Spearman: ρ = {spearman_r_old:.4f}, ρ² =
{spearman_r_old**2:.4f}")
        print(f"      Kendall:  τ = {kendall_tau_old:.4f}")

        results.append({
            'Condition': sheet_name,
            'Pole': 'Old',
            'n_cells': len(old_pole_mrfp),
            'Pearson_r': pearson_r_old,
            'Pearson_r_squared': pearson_r_old ** 2,
            'Pearson_p': pearson_p_old,
            'Spearman_rho': spearman_r_old,
            'Spearman_rho_squared': spearman_r_old ** 2,
            'Spearman_p': spearman_p_old,
            'Kendall_tau': kendall_tau_old,
            'Kendall_p': kendall_p_old
        })
    else:
        print(f"   ❌ OLD POLE: Not enough data ({len(old_pole_mrfp)}
cells)")

    #
======================================================================
==
    # NEW POLE (Pole 2)
    #
======================================================================
==

    new_pole_mrfp = df[NEW_POLE_MRFP].values
    new_pole_hada = df[NEW_POLE_HADA].values

    # Remove NaN
    mask_new = ~(np.isnan(new_pole_mrfp) | np.isnan(new_pole_hada))
    new_pole_mrfp = new_pole_mrfp[mask_new]
    new_pole_hada = new_pole_hada[mask_new]

    if len(new_pole_mrfp) >= 3:
        # Pearson
        pearson_r_new, pearson_p_new = stats.pearsonr(new_pole_hada,
new_pole_mrfp)

        # Spearman
        spearman_r_new, spearman_p_new =
stats.spearmanr(new_pole_hada, new_pole_mrfp)

        # Kendall
        kendall_tau_new, kendall_p_new =
stats.kendalltau(new_pole_hada, new_pole_mrfp)
```

```
        print(f"    📊 NEW POLE:")
        print(f"      n = {len(new_pole_mrfp)}")
        print(f"      Pearson:  r = {pearson_r_new:.4f}, r² =
{pearson_r_new**2:.4f}")
        print(f"      Spearman: ρ = {spearman_r_new:.4f}, ρ² =
{spearman_r_new**2:.4f}")
        print(f"      Kendall:  τ = {kendall_tau_new:.4f}")

        results.append({
            'Condition': sheet_name,
            'Pole': 'New',
            'n_cells': len(new_pole_mrfp),
            'Pearson_r': pearson_r_new,
            'Pearson_r_squared': pearson_r_new ** 2,
            'Pearson_p': pearson_p_new,
            'Spearman_rho': spearman_r_new,
            'Spearman_rho_squared': spearman_r_new ** 2,
            'Spearman_p': spearman_p_new,
            'Kendall_tau': kendall_tau_new,
            'Kendall_p': kendall_p_new
        })
    else:
        print(f"    ❌ NEW POLE: Not enough data ({len(new_pole_mrfp)}
cells)")

    print()

#
======================================================================
======
# SAVE RESULTS
#
======================================================================
======

if len(results) == 0:
    print("❌ No data was successfully analyzed!")
    exit(1)

results_df = pd.DataFrame(results)

# Save to CSV
output_file = 'batch_polar_correlations.csv'
results_df.to_csv(output_file, index=False)

print("=" * 80)
print("RESULTS SUMMARY")
print("=" * 80)
print()
```

```
print(results_df[['Condition', 'Pole', 'n_cells', 'Pearson_r_squared',
                  'Spearman_rho',
'Spearman_rho_squared']].to_string(index=False))

print("\n" + "=" * 80)
print("SAVED TO: " + output_file)
print("=" * 80)

#
======================================================================
======
# CREATE FORMATTED OUTPUT FOR PAPER
#
======================================================================
======

print("\n" + "=" * 80)
print("FOR YOUR PAPER (copy-paste ready)")
print("=" * 80)

for condition in results_df['Condition'].unique():
    cond_data = results_df[results_df['Condition'] == condition]

    print(f"\n{condition}:")

    for _, row in cond_data.iterrows():
        pole = row['Pole']
        rho = row['Spearman_rho']
        p = row['Spearman_p']
        n = int(row['n_cells'])
        p_str = f"p = {p:.2e}" if p >= 0.0001 else "p < 0.0001"

        print(f"  {pole} pole: ρ = {rho:.3f}, {p_str}, n = {n}")

print("\n✅ ANALYSIS COMPLETE!")
print(f"✅ Successfully analyzed {len(results_df)//2} conditions × 2
poles = {len(results_df)} datasets")
print(f"✅ Results saved to: {output_file}")
```

**Supplemental Figure 5. Batch septal correlation calculation python code.**

```
#!/usr/bin/env python3
"""
Calculates Pearson, Spearman, and Kendall correlations for
multiple datasets

HOW TO USE:
1. Put all your Excel files in the same folder as this script
2. Edit the FILES_TO_ANALYZE list below (lines 18-26)
3. Run: python batch_correlations.py
4. Results will be saved to: batch_correlation_results.csv

REQUIREMENTS:
- Excel files must have columns containing 'X' or 'Position' (for cell
position 0-1)
- Excel files must have columns with 'HADA' and 'Y' (for HADA
intensity)
- Excel files must have columns with 'pgfa' or 'mRFP' and 'Y' (for
mRFP intensity)
"""

import pandas as pd
import numpy as np
from scipy import stats
import os

#
======================================================================
======
# EDIT THIS SECTION - ADD OR REMOVE FILES AS NEEDED
#
======================================================================
======

FILES_TO_ANALYZE = [
    # Format: ('Display Name', 'filename.xlsx')
    ('Control', 'ctrl_both.xlsx'),
    ('Experimental', 'experimental.xlsx'),
]

# Optional: Change septal region boundaries (default: 0.4-0.6)
SEPTAL_START = 0.4
SEPTAL_END = 0.6

#
======================================================================
======
# NO NEED TO EDIT BELOW THIS LINE
#
======================================================================
```

```
======

def find_column(df, keywords):
    """Find column matching any of the keywords (case insensitive)"""
    for col in df.columns:
        col_lower = col.lower()
        if all(kw.lower() in col_lower for kw in keywords):
            return col
    return None

def analyze_file(filename, display_name):
    """Analyze one file and return correlation statistics"""

    try:
        # Check if file exists
        if not os.path.exists(filename):
            print(f"   ❌ File not found: {filename}")
            return None

        # Read Excel file
        df = pd.read_excel(filename)

        # Find required columns. Change 'pgfa' to 'your gene name' or
'your fp'. Change in
        # subsequent sections to prevent errors
        pos_col = find_column(df, ['x']) or find_column(df,
['position'])
        hada_col = find_column(df, ['y', 'hada'])
        mrfp_col = find_column(df, ['y', 'pgfa']) or find_column(df,
['y', 'mrfp'])

        if not all([pos_col, hada_col, mrfp_col]):
            print(f"   ❌ Could not find required columns")
            print(f"      Available columns: {list(df.columns)}")
            return None

        print(f"   📁 Using: Position={pos_col}, HADA={hada_col},
mRFP={mrfp_col}")

        # Filter for septal region
        septal = df[(df[pos_col] >= SEPTAL_START) & (df[pos_col] <=
SEPTAL_END)]

        if len(septal) == 0:
            print(f"   ❌ No data in septal region ({SEPTAL_START}-
{SEPTAL_END})")
            return None

        # Extract intensities
        hada = septal[hada_col].values
```

```
        mrfp = septal[mrfp_col].values

        # Remove NaN
        mask = ~(np.isnan(hada) | np.isnan(mrfp))
        hada = hada[mask]
        mrfp = mrfp[mask]

        if len(hada) < 3:
            print(f"   ❌ Too few valid datapoints ({len(hada)})")
            return None

        print(f"   ✅ {len(hada)} valid septal datapoints")

        # Calculate correlations
        pearson_r, pearson_p = stats.pearsonr(hada, mrfp)
        spearman_r, spearman_p = stats.spearmanr(hada, mrfp)
        kendall_tau, kendall_p = stats.kendalltau(hada, mrfp)

        # Display results
        print(f"   📊 Pearson:  r = {pearson_r:.4f}, r² =
{pearson_r**2:.4f}")
        print(f"   📊 Spearman: ρ = {spearman_r:.4f}, ρ² =
{spearman_r**2:.4f}")
        print(f"   📊 Kendall:  τ = {kendall_tau:.4f}")

        return {
            'Condition': display_name,
            'Filename': filename,
            'n_datapoints': len(hada),
            'Pearson_r': pearson_r,
            'Pearson_r_squared': pearson_r ** 2,
            'Pearson_p': pearson_p,
            'Spearman_rho': spearman_r,
            'Spearman_rho_squared': spearman_r ** 2,
            'Spearman_p': spearman_p,
            'Kendall_tau': kendall_tau,
            'Kendall_p': kendall_p
        }

    except Exception as e:
        print(f"   ❌ Error analyzing {filename}: {str(e)}")
        return None

#
======================================================================
======
# MAIN ANALYSIS
#
======================================================================
======
```

```
print("=" * 80)
print("BATCH SPEARMAN CORRELATION ANALYSIS")
print("=" * 80)
print(f"\nAnalyzing septal region: {SEPTAL_START}-{SEPTAL_END}")
print(f"Number of files to analyze: {len(FILES_TO_ANALYZE)}")
print()

results = []

for display_name, filename in FILES_TO_ANALYZE:
    print(f"Processing: {display_name}")
    result = analyze_file(filename, display_name)
    if result is not None:
        results.append(result)
    print()

#
======================================================================
======
# SAVE RESULTS
#
======================================================================
======

if len(results) == 0:
    print("❌ No files were successfully analyzed!")
    print("Check that:")
    print("  1. Files exist in the same folder as this script")
    print("  2. Filenames in FILES_TO_ANALYZE are correct")
    print("  3. Files have the required columns (X, Y-HADA, Y-pgfa)")
    exit(1)

results_df = pd.DataFrame(results)

# Save to CSV
output_file = 'batch_correlation_results.csv'
results_df.to_csv(output_file, index=False)

print("=" * 80)
print("RESULTS SUMMARY")
print("=" * 80)
print()
print(results_df[['Condition', 'n_datapoints', 'Pearson_r_squared',
                  'Spearman_rho',
'Spearman_rho_squared']].to_string(index=False))

print("\n" + "=" * 80)
print("SAVED TO: " + output_file)
print("=" * 80)
```

```
# Create a summary for easy copy-paste into report
print("\n" + "=" * 80)
print("FOR YOUR REPORT (copy-paste ready)")
print("=" * 80)

print("\nSpearman correlation coefficients (septal region):")
for _, row in results_df.iterrows():
    rho = row['Spearman_rho']
    p = row['Spearman_p']
    n = int(row['n_datapoints'])
    p_str = f"p = {p:.2e}" if p >= 0.0001 else "p < 0.0001"
    print(f"  {row['Condition']:12s}: ρ = {rho:.3f}, {p_str}, n =
{n}")

print("\n ANALYSIS COMPLETE!")
print(f"Successfully analyzed {len(results)}/{len(FILES_TO_ANALYZE)}
files")
print(f"Results saved to: {output_file}")
```
